# Nuclear Transformation In Metalloenzyme. A Novel And High Potential Cancer Treatment Research

**DOI:** 10.1101/2023.08.10.552823

**Authors:** Tran Van Luyen, Truong Hoang Tuan

**Affiliations:** Nuclear Research Institute. Dalat Vietnam; Center for Nuclear Techniques, HoChiMinh city, Vietnam

## Abstract

In this study, we have introduced a novel approach to cancer treatment involving the deactivation of metalloenzymes through the utilization of radioisotopes. The concept of leveraging radioisotopes to interact with metalloenzymes represents a groundbreaking theoretical advancement. Through simulations utilizing the MIRD code and based on the consistent concentration of stable Mg within stage 2A cancerous tissue, we have quantified the potential success rates.

To conduct these simulations, we employed 0.1 nanograms (ng) of stable Mg, which corresponds to an activity of 19.7 MBq of Mg-28. This data was input into the MIRD calculations to estimate the absorbed doses within various organs, employing diverse methods of radioisotope administration into the body. Remarkably, even with a mere 1‰ probability of effectively reaching the intended cancerous tissues, this quantity of Mg-28 demonstrates the capability to render billions of Mg-containing metalloenzymes inactive.

The remarkable efficiency achieved through precise radioisotope targeting underscores the promise of this methodology. Nevertheless, the findings underscore the necessity of undertaking both in vitro and in vivo research initiatives prior to embarking on clinical trials.

## Overview

Enzyme inactivation methods have found utility in biotechnology and cancer chemotherapy [1-5]. These techniques often involve the use of natural or artificial complexes to hinder pure protein enzymes or apoenzymes within metalloenzymes or coenzymes. However, the quest for suitable inhibitors capable of complete enzyme inactivation without inducing adverse effects remains a formidable challenge. Thus far, no single compound, complex, or combination of inhibitors has delivered the anticipated outcomes. A solution to this intricate problem might lie in a simpler and more effective approach—specifically, the notion of utilizing radioisotope isotopes to inactivate metalloenzymes.

Metalloenzymes comprise two integral constituents: the apoenzyme and the pivotal metal ion cofactor, wherein the latter exerts a critical influence on the enzyme’s catalytic attributes and functions. Both constituents exhibit robust stability with regard to complex formation and binding. In terms of biochemical bonding, cofactors exhibit a distinctive nature that resists replacement by alternative metal ions via conventional chemical methodologies. Perturbations to these cofactor ions or complexes lead to the inactivation of metalloenzymes. While substituting the apoenzyme proves ineffective, as mentioned earlier, a novel strategy involves the substitution of metal ions as a means of inactivation. This approach hinges on answering two fundamental questions: i) Can the current metal ions be substituted with alternate metal ions? and ii) If so, what strategies can effectuate this substitution? Addressing these inquiries holds the potential to definitively resolve the conundrum of metalloenzyme inactivation.

The spectrum of metalloenzymes in Homo sapiens is extensive [6], and several of these enzymes are integral to DNA replication, protein synthesis, and ATP metabolism—critical processes in cellular cycles. This study is centered on three such enzymes: Hexokinase, DNA polymerase (DNA-pol), and RNA polymerase (RNA-pol). Hexokinase propels the ATP → ADP reaction, yielding energy [7], and operates within the cell’s cytoplasm. DNA-pol spearheads DNA replication [8] and is situated within the cell nucleus amidst chromosomes. RNA-pol catalyzes processes involved in diverse protein creation [9]. Each of these enzymes relies on Mg2+ ions as cofactors. Inactivating these enzymes becomes conceivable by replacing the Mg ions. Such a strategy could extend to cofactors housing distinct metal ions.

Magnesium (Mg) assumes vital roles within the body, serving not only as a protein constituent but also as a cofactor for a myriad of enzymes, numbering over 300 distinct types [10-12]. In its natural state, magnesium exhibits three stable isotopes: Mg-24 (0.79%), Mg-25 (0.10%), and Mg-26 (0.11%) [13]. Intriguingly, neither natural nor synthetic compounds can distinguish between Mg-24, Mg-25, or Mg-26 isotopes when bearing Mg2+ ions. This characteristic presents the possibility of substituting stable Mg ions with specific radioisotope isotopes of magnesium, while retaining the chemical attributes of these compounds.

Upon the decay of a radioisotope isotope of magnesium, it metamorphoses into an element either directly preceding or following magnesium within the periodic table. The viability of this transformation depends on the mode of decay. Mg-28, possessing a half-life of roughly 21 hours, emerges as an optimal candidate for replacing stable Mg ions within enzymes. Its prolonged decay period facilitates both experimental and medical procedures. Mg-28 experiences beta (electron) decay, generating Al-28, which subsequently undergoes further beta decay to attain stable Si. Notably, neither Al3+ nor Si4+ ions serve as cofactors for the mentioned enzymes, leading to their instantaneous inactivation [14-17]. However, the essential cellular metabolic processes dependent on these enzymes necessitate continuity. To address the absence of catalyzers for biochemical reactions, cells promptly initiate the production of new enzymes to substitute the inactivated ones. This pattern persists as the freshly formed enzymes similarly succumb to inactivation, compelling cells to perpetuate enzyme regeneration. This cycle ceases only upon the full decay of the radioisotope isotopes of Mg-28 or their scarcity relative to the omnipresent stable Mg ions, which consistently sustain cellular nourishment through the circulatory system. When this mechanism is solely targeted at cancer cells, it emerges as a novel cancer treatment approach, leveraging the exclusive inactivation of metalloenzymes through the radioisotope isotope Mg-28. However, a challenge arises in that this method not only incapacitates enzymes in cancerous cells but also impinges on those in healthy cells. Concurrently, the radiation effect manifests.

Radiation’s impact on tissue cells directly impairs DNA molecules and other proteins within the targeted cells. Additionally, radiation engenders a cascade of free radicals along its trajectory. The phenomenon produces ions such as Mg, Al, and Si through nuclear decay reactions, these ions either recoiling or forming. Subsequent collisions with protein molecules in the intracellular fluid are feasible, with the recoiling energy deemed sufficient for severing chemical bonds in their path. In terms of physics, the application of this method in cancer treatment demands the consideration of an effective safe radiation dosage. Through utilization of the Mg-28 isotope’s Sv/Bq conversion factor in ICRP 119 [18], along with mass and tissue weighting factors elucidated in ICRP 53 [19], dose calculation formulations outlined in ICRP 103 [20], and guided by MIRD calc algorithms [21] underpinned by nuclear data from ICRP 107 [22], computations will ascertain the effective whole-body dose and tissue-specific doses stemming from 19.7 MBq (0.53 mCi)—equivalent to 0.1 ng of Mg-28. Further analysis will unveil dose distributions contingent upon the route of Mg-28 incorporation into the body.

The Mg-28 isotope exhibits a maximum beta radiation energy capable of penetrating water to a depth of 0.36 cm [23]. Given that our bodies consist of approximately 70%-75% water, this range can be interpreted as the utmost distance beta radiation can traverse within tissue. Should radioactive isotopes be situated at the tumor mass periphery, the effective radius of action will mimic this shape, enveloping the tumor mass with dimensions equivalent to the sum of the tumor’s size and an additional 0.36 cm. However, this scenario yields an undesirable outcome by affecting neighboring normal tissues surrounding the cancerous mass. The total energy of beta particles emitted from the Mg-28 isotope is Qβ=1830.77 KeV [23], and with a maximum range of 0.36 cm, the calculation yields an average linear energy transfer (LET) of beta particles from Mg-28 at 0.508 KeV/μm or 0.508 eV/nm.

Mg-28 has been employed since its discovery in 1953 [23] for metabolic investigations in plants [24-26], followed by analogous studies on animals’ physiology and metabolism for several decades [27-38]. The intrigue surrounding Mg-28’s application for studying Mg’s pathology, absorption, excretion, and metabolic dynamics within the human body has also captured researchers’ attention [39-47]. However, to date, no explorations have scrutinized Mg’s role within enzymatic cofactors using the radioactive Mg-28 isotope. Furthermore, no inquiries have tackled the inactivation of these enzymatic cofactors using Mg-28.

In the context of crucial Mg-containing enzymatic cofactors like DNA pol and RNA pol, numerous investigations have ventured into the connection between Mg and carcinogens [48], explored DNA pol γ and DNA pol π [49], and probed the interaction of Mg with enzymes [50-57]. However, these inquiries have refrained from utilizing the radioactive tracer form of Mg-28 and have not ventured into the inactivation of these enzymes via Mg-28.

Historically, cancer treatment methods encompass surgery, chemotherapy, radiotherapy, immune modulation, thermal laser, and tumor vascular targeting [58,59]. The National Cancer Institute (NCI), a branch of the NIH [60], as well as Dr. M.R.A. Pillai’s doctoral thesis [61], comprehensively elucidate these methods. Mention is made of targeted cancer therapy utilizing small molecules to effectively access and infiltrate cancer cells for damage infliction [60]. While the medical applications of radioactive isotopes for diagnosis and therapy are well-established, our knowledge suggests a paucity of investigations into the use of the radioactive tracer form of Mg-28 for targeted cancer therapy.

The technique of enzyme inactivation using radioactive isotopes, though presently lacking empirical data, harbors the potential to deliver radioactive isotopes directly to the enzyme molecules engaged in metabolic activities. Essentially, radiation would be brought to the target site and subsequently decay. In effect, this approach accomplishes two pivotal objectives in cancer treatment: i) impeding vital metabolic processes within cancerous cells, notably those governing energy provision, DNA replication, and the synthesis of characteristic proteins throughout the cellular cycle, and ii) exterminating cancer cells by means of ionizing radiation at low dosages while ensuring an unequivocal target strike probability of 100%.

The merit of this method resides in the fusion of classical strategies like chemotherapy to suppress essential metabolic functions and the precision of targeted radiation therapy, thereby refining and augmenting their efficacy.

Nevertheless, further investigation is imperative to address the logistical aspect of delivering radioactive isotopes as cofactors to the tumor site, as well as to understand the competitive interplay between radioactive ions and stable ions within the enzymes they serve as cofactors. This scenario mirrors the case of I-131’s competition with stable iodine within the thyroid gland structure during thyroid cancer treatment [64].

### II. Research Method

#### II.1. Theoretical Basis

Metalloenzymes have a structure consisting of a Protein + metal ion.

We denote the Protein as P, the metal ion as M, and M* as the radioactive isotope of the metal [14]. The symbols *(s)* correspond to the stable state, *(i)* represents the inhibited state, and *(d)* represents the decay state of the radioactive isotope. We have the following possible states of the metalloenzyme:

Thus, to prevent the metabolic process within the cell through the metalloenzyme, we have three approaches:

a. Inhibit the Protein.
b. Replace the metal ion with its radioactive isotope.
c. Both inhibit the Protein and use the radioactive isotope as a substitution for the metal ion.

We choose the second approach (b), replacing the stable metal ion with its appropriate radioactive isotope. This method has been elucidated in previous studies [14,15].

#### II.2. Analysis of Mg Content in Cancerous and Healthy Tissue Samples

Numerous methods for elemental analysis possess the requisite sensitivity to determine magnesium (Mg) content in biological samples, especially those of medical significance [64, 65]. Among these methods, atomic absorption spectroscopy has been chosen [66]. The acquisition and examination of medical samples adhere to stringent standards, necessitating approval from the medical ethics council. The Oncology Hospital of Ho Chi Minh City Medical Council granted approval for the utilization of cervical and colorectal cancer tissues. Table 2 provides the sample weights and codes. Employing farafin packaging, the samples were dispatched to the Dalat Nuclear Research Institute for Mg analysis via atomic absorption spectroscopy. The findings of this analysis are tabulated in Table 1. To enhance clarity, the results are presented in the unit of 10^-6g/g (parts per million, ppm).

**Table 1.** States of metalloenzyme.

| No. | Protein | Metal Ion | Metalloenzyme | Status of enzymes |
| --- | --- | --- | --- | --- |
| 1 | $P_s$ | $M_s$ | Metalloenzyme | Active |
| 2 | $P_i$ | $M_s$ | Non Metalloenzyme | Inhibition |
| 3 | $P_s$ | $M^*_d$ | Non Metalloenzyme | Inactivate |
| 4 | $P_i$ | $M^*_d$ | Non Metalloenzyme | Damage |

**Table 2.** Magnesium ion content in the analyzed samples, magnesium content in stage 2A samples, and the equivalent activity of Mg-28 for effective dose calculation in tissues.

| Sample Code | Tissue Sample Name | Result (mg/kg) | Mg content in stage 2a (ng) | Equivalent activity of Mg-28 (MBq) |
| --- | --- | --- | --- | --- |
| DV (22) 4199 A' | Breast Cancer | 16,11 ± 1,21 | 0,11 ± 0,01 | 22,22 ± 0,32 |
| DV (22) 4199 A | Breast Cancer | 14,52 ± 1,12 | 0,10 ± 0,01 | 19,72 ± 0,51 |
| DV (22) 4194 A' | Breast Cancer | 14,31 ± 1,32 | 0,09 ± 0,01 | 17,73 ± 0,22 |
| DV (22) 4194 A | Breast Cancer | 11,81 ± 1,04 | 0,08 ± 0,01 | 15,73 ± 0,23 |
| DV (22) 4199-L' | Healthy Breast Tissue | 16,92 ± 1,51 | <u>0,12 ± 0,04</u> | <u>23,62 ± 0,31</u> |
| DV (22) 4199-L | Healthy Breast Tissue | 7,62 ± 3,52 | <u>0,05 ± 0,01</u> | <u>9,82 ± 0,54</u> |
| DV (22) 4161-21A' | Cervix Uteri | 54,53 ± 3,81 | 0,38 ± 0,05 | 78,42 ± 0,72 |
| DV (22) 4161-21A | Cervix Uteri | 50,94 ± 3,52 | 0,35 ± 0,03 | 68,92 ± 0,64 |
| DV (22) 4648 -LNC' | Healthy Colon | 38,62 ± 3,03 | <u>0,27 ± 0,02</u> | <u>53,14 ± 0,52</u> |
| DV (22) 4648-LNC | Healthy Colon | 43,03 ± 3,31 | <u>0,30 ± 0,03</u> | <u>59,14 ± 0,43</u> |
| DV (22) 4648-ANC' | Colon Cancer | 41,72 ± 4,23 | 0,29 ± 0,04 | 57,15 ± 0,61 |
| DV (22) 4648-ANC | Colon Cancer | 55,71 ± 5,13 | 0,39 ± 0,04 | 74,82 ± 0,72 |

For calculations pertaining to the 2A stage tumor, it is postulated that the tumor possesses dimensions of 2x3x5 cm. This assumption aids in evaluating the aggregate Mg content within it. Notably, the overall Mg content within the 2A stage tumor ranges from 0.36 ng to 1.72 ng. Utilizing 0.1 ng of Mg as a basis, the formula:

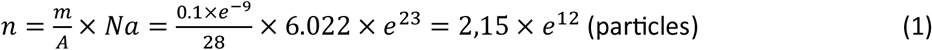

is employed, where *m* signifies the Mg amount (0.1 ng), *A* represents the atomic weight of Mg-28 (28), *Na* denotes Avogadro’s number, and *n* symbolizes the number of Mg ion particles. Assuming all these ions are Mg-28 isotopes and considering a 1‰ likelihood of interaction with the tumor, the outcome is an estimated 2.15 x 10^9 enzymes that can be inactivated. This translates to an activity of 19.7 MBq (0.53 mCi). The calculation for activity is expressed by:

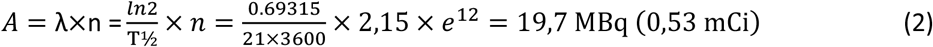

Here, *λ* signifies the decay constant of Mg-28, *n* represents the particle count calculated in equation (1), and *T*_1/2_ denotes the half-life of Mg-28, which stands at 21 hours. Thus, the value of 0.1 ng of Mg-28 serves as the basis for subsequent calculations using the MIRD program [21].

#### II.3. Calculation of Mg-28 Absorbed Dose. Calculation Methods using MIRD

MIRD [21] is an acronym that stands for “Medical Internal Radiation Dose.” It represents a mathematical framework employed for estimating the radiation dose absorbed by organs and tissues within the human body due to internal sources of radioactivity. This model finds extensive use in nuclear medicine with the primary goal of optimizing the benefits of a particular radiation source for diagnostic or therapeutic purposes while minimizing any potential unintended side effects. The foundation of the MIRD model rests on several fundamental assumptions. These encompass anatomical and kinetic models of the absorption, distribution, metabolism, and excretion of radioactive isotopes within the body. This, in turn, incorporates nuclear properties intrinsic to the radiation source, including its type, energy, and half-life.

MIRD undertakes calculations to determine absorbed dose and effective dose, shaping radiation therapy protocols, evaluating the risks tied to radiation exposure, and gauging the safety and effectiveness of novel radiopharmaceuticals. Essentially, the MIRD model constitutes a pivotal tool in optimizing the judicious utilization of radiation in medical applications, and its continuous refinement and updates bolster its significance.

The application of this method involves computing the effective dose for various isotopes, encompassing I-131, Y-90, Lu-177, and with particular emphasis, Mg-28—representing the central focus of this study. Distinct input data scenarios, encompassing intravenous injection, oral ingestion, and inhalation, were simulated to discern the absorbed dose corresponding to 19.7 MBq (0.53 mCi) associated with 0.1 ng of Mg-28. These calculations took into account different source and target tissues, notably including tumor masses (designated as tissues 1-5). The comprehensive outcome of these computations is presented in tables 3 through 6.

**Table 3.**
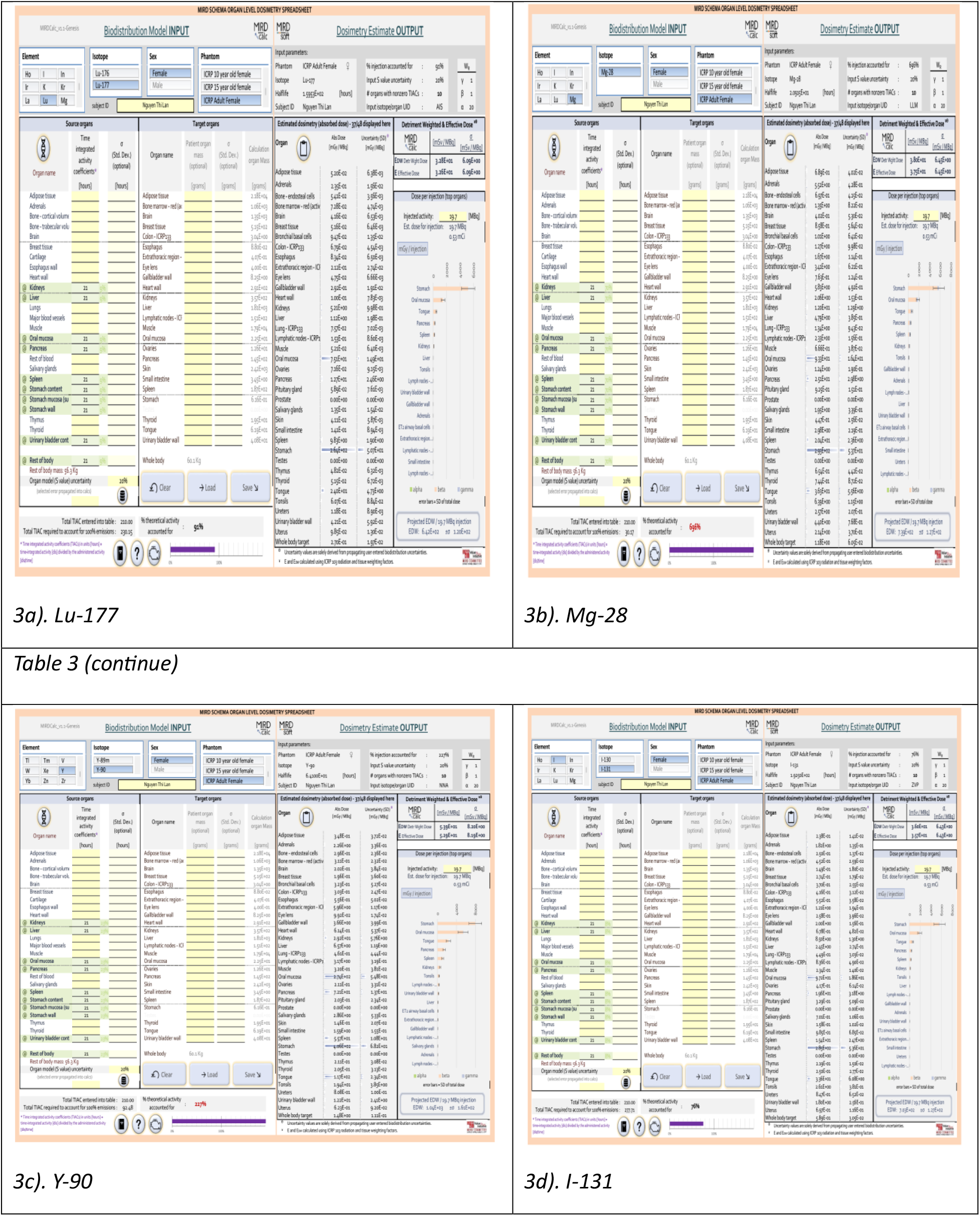
MIRD calculations results for oral ingestion through the stomach with isotopes Lu-177 (3a), Mg-28 (3b), Y-90 (3c), and I-131 (3d) on the same phantom subject and at the same activity level of 19.7 MBq, with retention over 21 hours.

### III. Results and Discussion

#### III.1. Results

The findings stemming from the analysis of magnesium content within both cancerous and healthy tissue samples of the same type are meticulously presented in Table 2. Within this table, we have meticulously computed magnesium content and the corresponding activity of Mg-28 within hypothesized stage 2a cancer tissue samples, each of a size ≤ 7cc. Table 3 is dedicated to showcasing simulated absorbed dose values linked to various radioactive isotopes frequently employed in cancer treatment—namely Lu-177, Y-90, I-131—alongside Mg-28. These calculations encompass the gastrointestinal route (via stomach, small intestine) as well as the respiratory route (inhalation), all at an identical activity level of 19.7 MBq. These discernments are concisely summarized in Table 4.

**Table 4.**
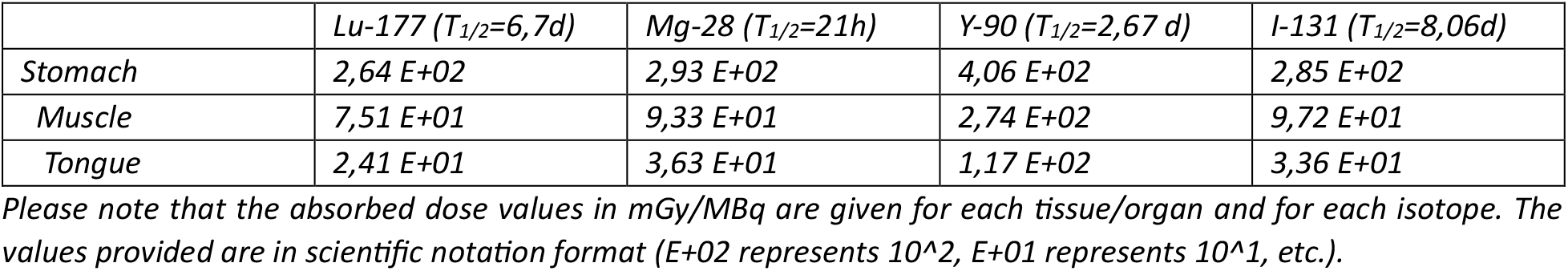
Absorbed dose per unit activity (mGy/MBq) for oral ingestion into the stomach of the four isotopes Lu-177, Mg-28, Y-90, and I-131.

Table 5 is dedicated to simulated absorbed dose values for 0.1 ng of Mg-28, presenting distinct avenues of entry into the body. On the other hand, Table 6 furnishes calculated absorbed dose values for 0.1 ng of Mg-28 following intravenous injection across various organs and tumor masses of varying dimensions, subsequent to a 21-hour interval. The nomenclature employed ranges from T1 to T4 for tumor masses not exceeding 7cc in size, and T5 to denote a 500cc mass.

**Table 5.**
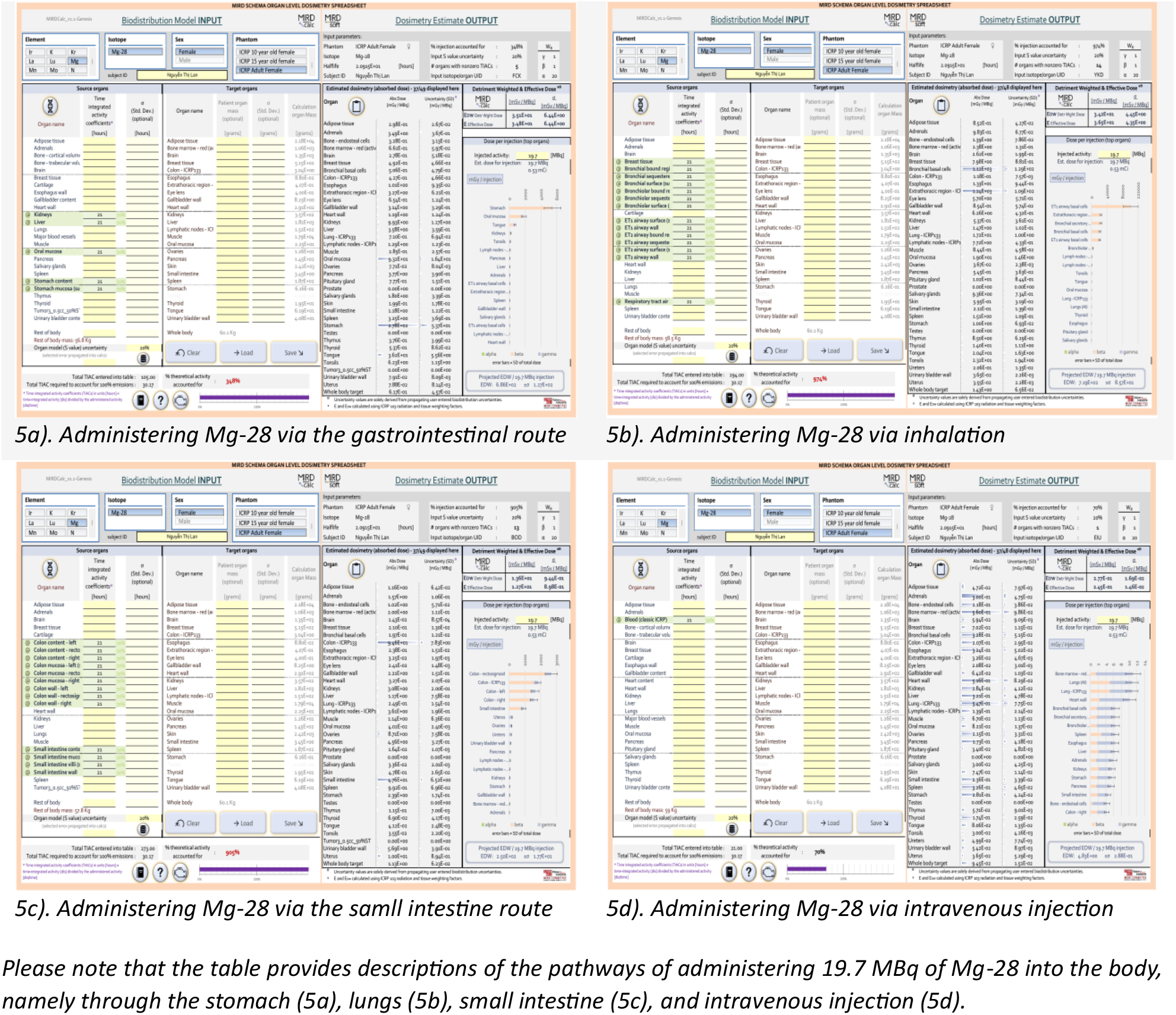
Pathways of administering 19.7 MBq of Mg-28 into the body with a retention time of one half-life (21 hours).

**Table 6.**
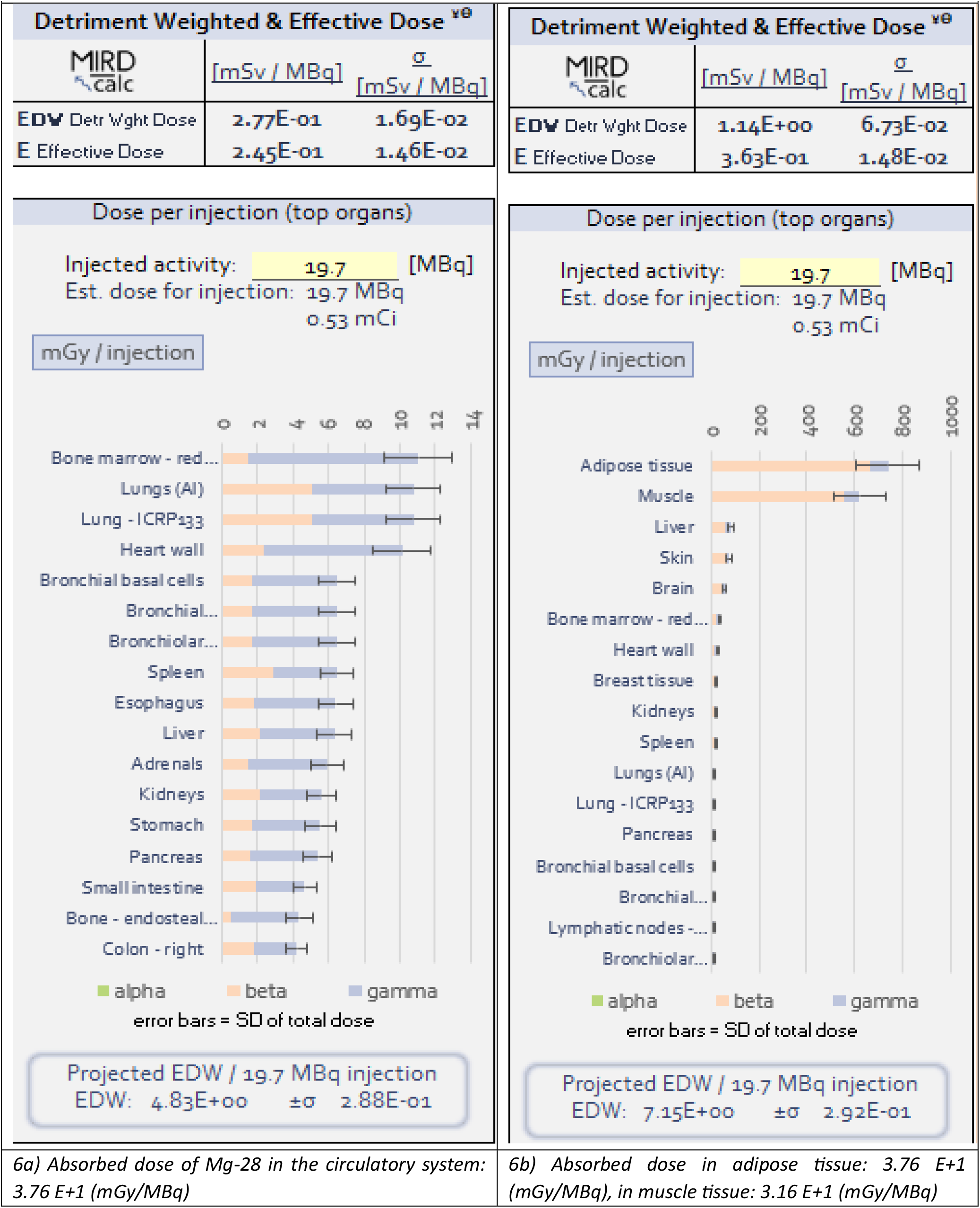
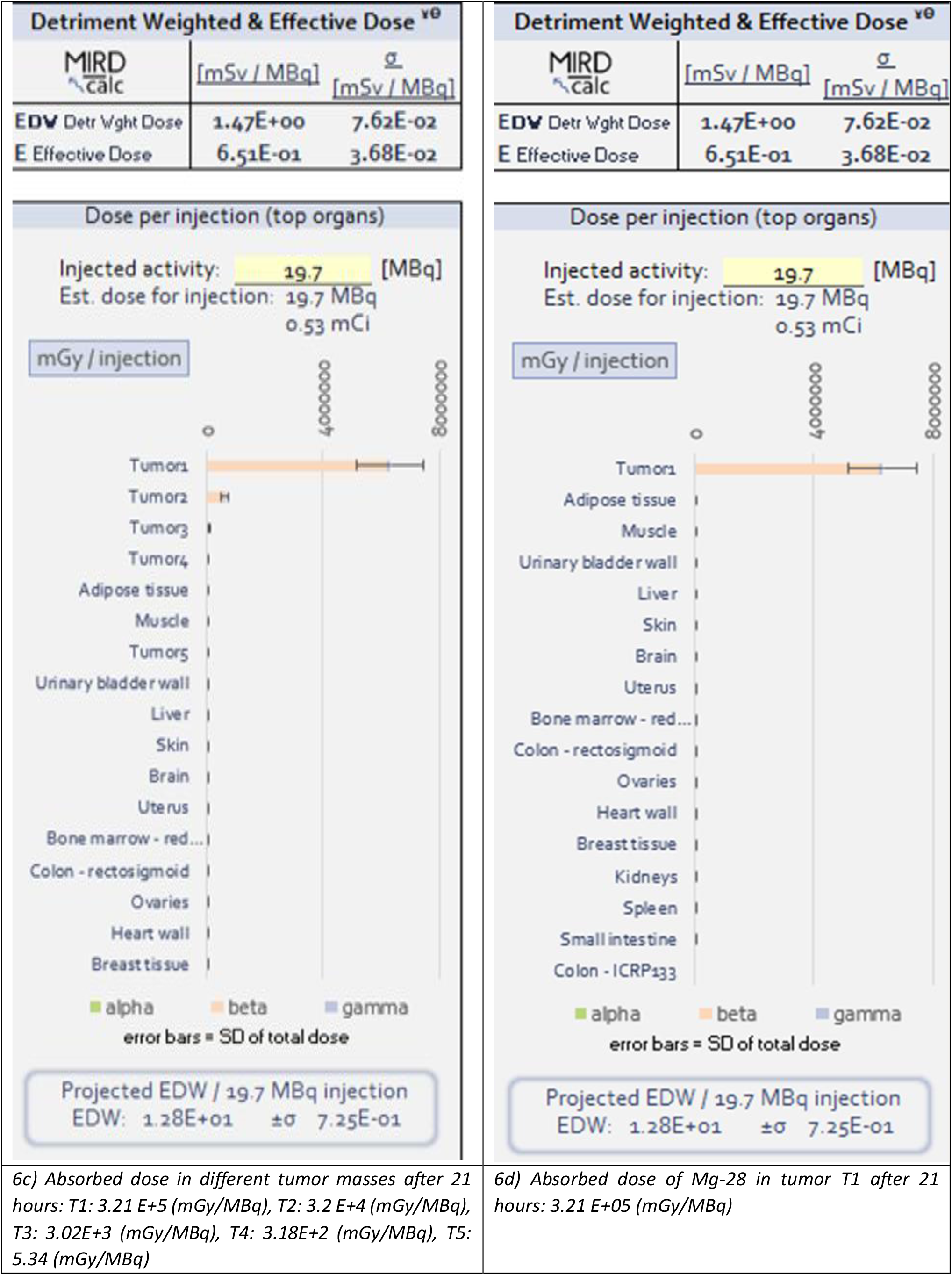
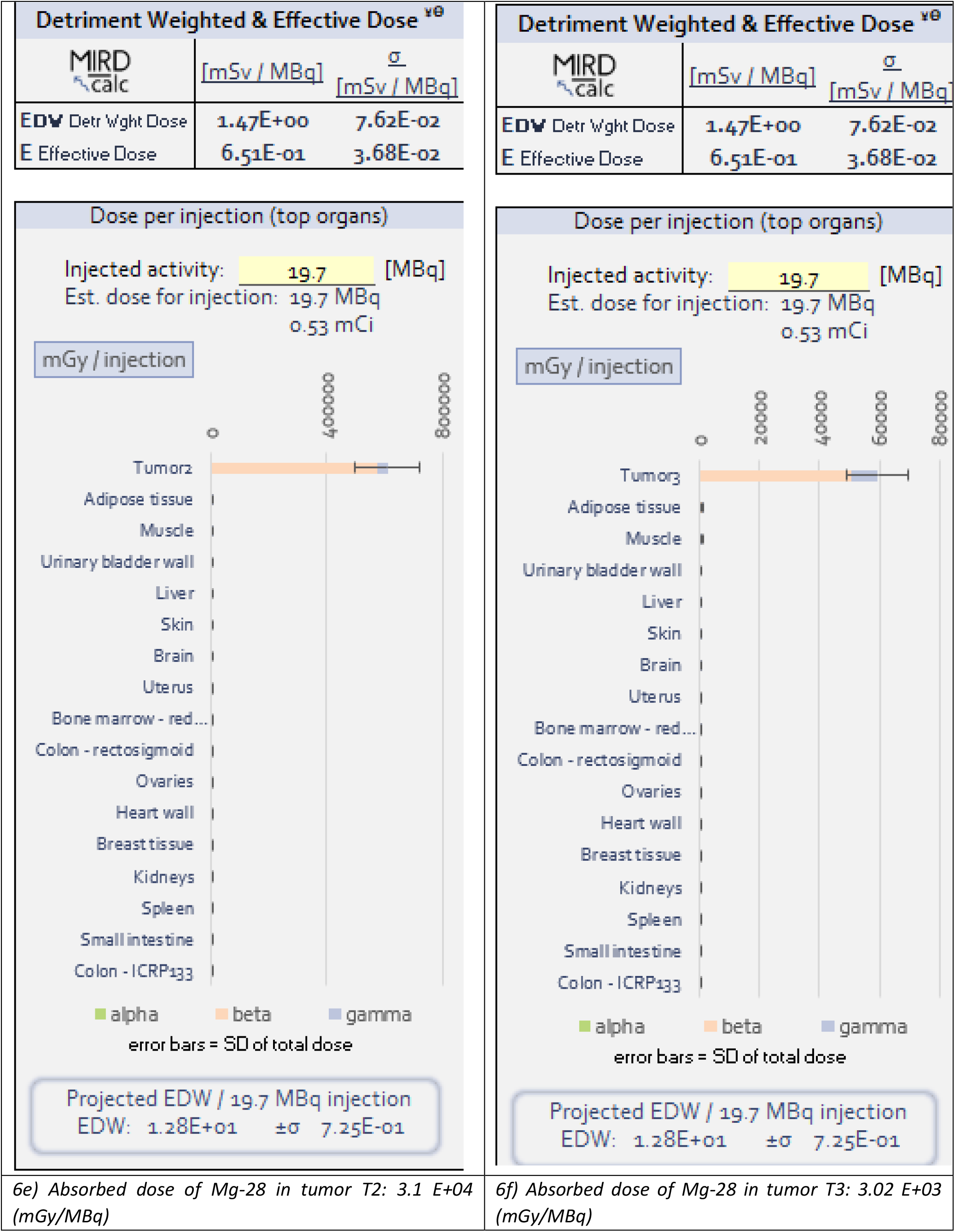
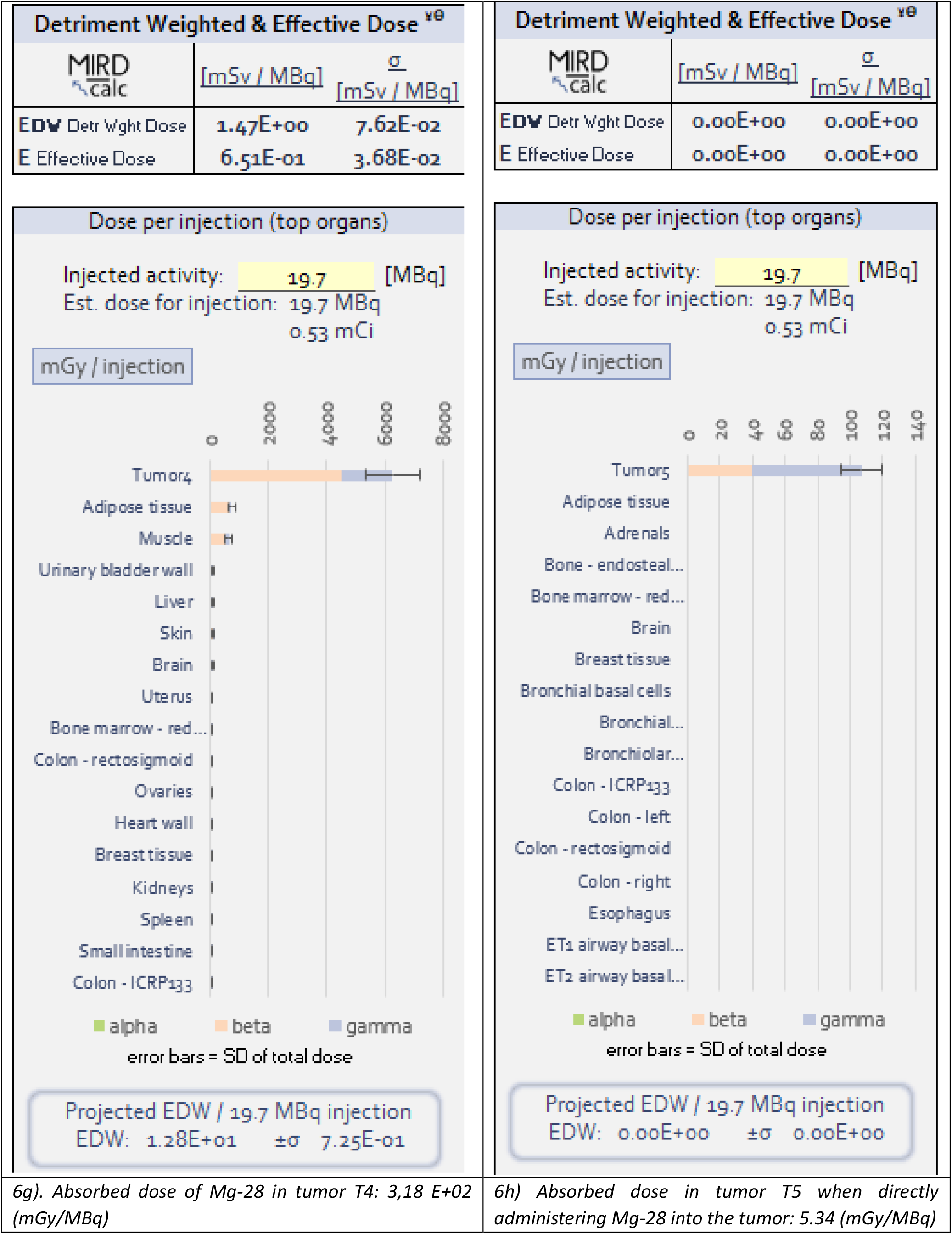
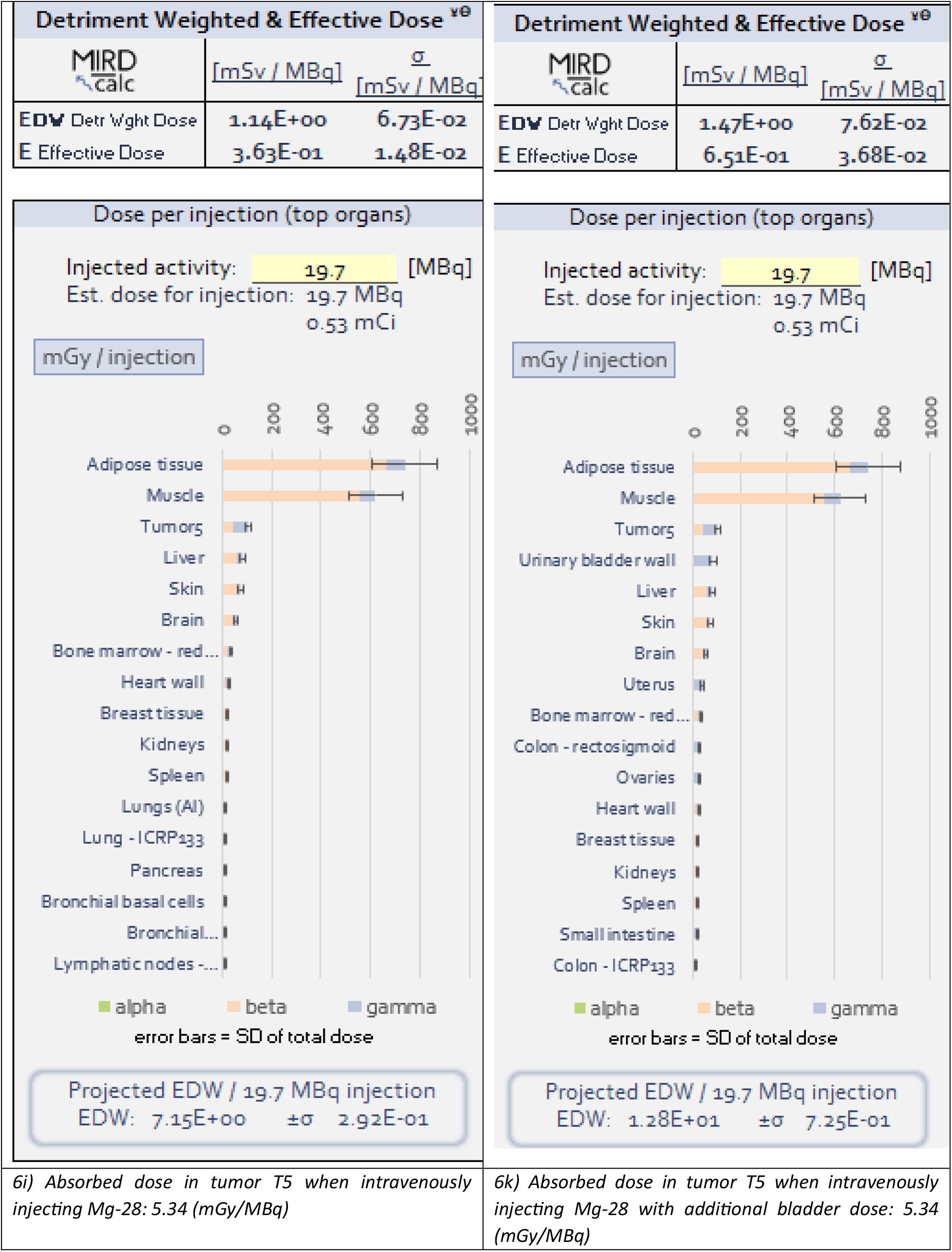
Absorbed dose of Mg-28 via intravenous injection in different tissues and U masses after 21 hours.

| Detriment Weighted & Effective Dose <sup>v8</sup> |  |  | Detriment Weighted & Effective Dose <sup>v8</sup> |  |  |
| --- | --- | --- | --- | --- | --- |
| MIRD<br>calc | [mSv / MBq] | $\sigma$<br>[mSv / MBq] | MIRD<br>calc | [mSv / MBq] | $\sigma$<br>[mSv / MBq] |
| EDW Detr Wght Dose | 2.77E-01 | 1.69E-02 | EDW Detr Wght Dose | 1.14E+00 | 6.73E-02 |
| E Effective Dose | 2.45E-01 | 1.46E-02 | E Effective Dose | 3.63E-01 | 1.48E-02 |

| Dose per injection (top organs) |  | Dose per injection (top organs) |
| --- | --- | --- |
| Injected activity: 19.7 [MBq]<br>Est. dose for injection: 19.7 MBq<br>0.53 mCi |  | Injected activity: 19.7 [MBq]<br>Est. dose for injection: 19.7 MBq<br>0.53 mCi |
| mGy / injection |  | mGy / injection |
| Projected EDW / 19.7 MBq injection<br>EDW: 4.83E+00 ±σ 2.88E-01 |  | Projected EDW / 19.7 MBq injection<br>EDW: 7.15E+00 ±σ 2.92E-01 |

|  |  |
| --- | --- |
| 6a) Absorbed dose of Mg-28 in the circulatory system:<br>3.76 E+1 (mGy/MBq) | 6b) Absorbed dose in adipose tissue: 3.76 E+1<br>(mGy/MBq), in muscle tissue: 3.16 E+1 (mGy/MBq) |

| Detriment Weighted & Effective Dose <sup>v0</sup> |  |  |
| --- | --- | --- |
| MIRD<br>calc | [mSv / MBq] | $\sigma$<br>[mSv / MBq] |
| EDW Detr Wght Dose | 1.47E+00 | 7.62E-02 |
| E Effective Dose | 6.51E-01 | 3.68E-02 |

| Detriment Weighted & Effective Dose <sup>v0</sup> |  |  |
| --- | --- | --- |
| MIRD<br>calc | [mSv / MBq] | $\sigma$<br>[mSv / MBq] |
| EDW Detr Wght Dose | 0.00E+00 | 0.00E+00 |
| E Effective Dose | 0.00E+00 | 0.00E+00 |

6h) Absorbed dose in tumor T5 when directly administering Mg-28 into the tumor: 5.34 (mGy/MBq)

According to the insights presented in Table 6, intravenous injection of Mg-28 without a concurrent tumor mass leads to widespread distribution across diverse organs, akin to scenario (5d). This scenario assumes the delivery of Mg-28 to organs via the circulatory system (6a), retention within the residual body portion (6b), and dose dispersion across varying tumor masses (6c), (6d), (6e), (6f), (6g), (6h). Notably, figures (6i) and (6k) provide a visualization of dose distribution within tumor mass T5, incorporating intravenous injection, with consideration for bladder retention and the residual body portion. Of particular significance, figure (6h) postulates a scenario where Mg-28 is directly introduced into tumor mass T5.

##### III.2. Discussion

Observing the data in column 3 of Table 2, it becomes evident that the magnesium content in tumor masses and healthy tissues shows minimal variance, aligning within the range of 7.62 to 16.92 ppm in breast tissue and 38.62 to 55.71 ppm in colon tissue. These values adhere consistently to established references [68-73]. Meanwhile, column 4 of the same table details the magnesium quantities in stage 2A tumors, and column 5 computes the corresponding Mg-28 activity. Hence, the magnesium ion content within the three categories of stage 2A tumors spans from 0.1 to 0.4 nanograms. Notably, assuming that 0.1 ng of Mg-28 can engage in competition with magnesium within cancer cells at an exceedingly low probability (1‰), it could potentially deactivate 2.15 x 10^9 enzymes reliant on magnesium ions as cofactors. It’s worth mentioning that highlighted values belonging to healthy tissues within the table can be disregarded. Additionally, the value of 0.1 ng of Mg-28, corresponding to 19.7 MBq or 0.53 mCi of radiation as deduced from equation (2), yields effective dose values suitable even for substantial tumor masses, while inflicting minimal impact on critical healthy tissues such as the heart, lungs, uterus, liver, kidneys, and spleen (as reflected in Tables 5 and 6).

A careful examination of Table 3 reveals the disposition of radioactive substances—Lu-177 (3a), Mg-28 (3b), Y-90 (3c), and I-131 (3d)—following oral administration after 21 hours (spanning a half-life of Mg-28 for comparison). These substances predominantly accumulate within the stomach. Disparities emerge solely in terms of dosage (MBq/mGy), with Y-90 displaying elevated doses in the stomach, muscle, and tongue compared to the remaining three isotopes. Corresponding reference values are outlined in Table 4.

Upon consulting Table 5, it’s evident that the distribution of Mg-28 activity within the body after 21 hours significantly varies based on the chosen administration pathways. Orally or inhalationally introduced Mg-28 primarily concentrates its radiation impact on tissues such as the stomach, small intestine, and lungs. This phenomenon is attributed mainly to beta radiation, with gamma radiation exerting minimal influence (scenarios 5a, 5b, and 5c). In contrast, the distribution pattern shifts when Mg-28 is introduced intravenously, leading to its relatively widespread distribution across diverse organs. The absorbed dose calculation encompasses contributions from both beta and gamma radiation (scenario 5d). Consequently, for tumors located within the digestive and respiratory organs, oral ingestion or inhalation could be the preferred administration routes. Conversely, for tumors in other internal organs, intravenous injection might be a more suitable choice.

Further insight can be gleaned from the scenario where Mg-28 is solely administered intravenously after 21 hours. In this instance, blood serves as the medium for transporting and dispersing Mg-28 to various organs, as demonstrated in Figure 6a. This distribution proves relatively comprehensive, with the absorbed dose being relatively modest, approximately 12 mGy per injection per organ. In scenarios where the residual body portion retains Mg-28, the distribution pattern aligns with Figure 6b, concentrating within adipose and muscle tissues. Here, the absorbed dose is even smaller, approximating 800 mGy per injection. As fat and muscle tissues constitute roughly 50-60% of body weight and boast a widespread presence, the concentration of Mg-28 within these tissues incurs minimal impact on crucial organs. Hence, this dose could be deemed safe for the body. Consequently, administering 0.1 ng of Mg-28 aligns with radiation exposure safety considerations.

Analyzing the absorbed dose distribution of Mg-28 within various tumor masses underscores its stability and a direct correlation with tumor size. The absorbed dose values are delineated as follows: U1: 3.21 x 10^5 mGy, U2: 3.2 x 10^4 mGy, U3: 3.02 x 10^3 mGy, U4: 3.18 x 10^2 mGy, U5: 5.34 mGy/MBq. Evidently, the smaller the tumor mass, the higher the absorbed dose, owing to the mass reduction.

Absorbed dose, as defined, is the radiation energy absorbed per unit mass of the material it traverses.

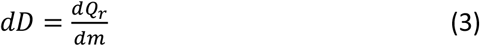

Here:

- dD is the absorbed dose calculated in Gy and has the dimension J/kg (Joules per Kilogram).
- dQr is the radiation energy of a decay, measured in eV or multiples of eV such as keV, MeV, or GeV. In the case of isotopes located inside, the energy loss from the body due to highly penetrating gamma or X-rays is negligible compared to the portion absorbed by high-energy charged particles. Therefore, dQr can be considered as the radiation energy absorbed. It should be noted that 1 eV = 1.6 x 10^-19 J.
- dm is the mass of the material through which the radiation passes, measured in kg.

Hence, it becomes evident that the smaller the tumor mass, the higher the absorbed dose, yielding a more pronounced interaction between radiation and the tumor mass. In the case of U5, direct delivery of Mg-28 to the tumor mass without intravenous injection sustains the absorbed dose at 5.34 mGy/MBq, mirroring the outcome of intravenous injection. However, notable differences arise in other organs such as adipose tissue, muscle tissue, and the bladder, where radiation retention is absent. This point underscores a crucial consideration: if a means of directly delivering Mg-28 to the tumor mass can be devised, potential side effects could be minimized.

Shifting focus to tumor masses T1 through T4, the absorbed dose of Mg-28 showcases considerable elevation, holding the potential to eliminate cancer cells with a single injection. The absorbed dose values per injection are outlined as follows: T4: 6000 mGy/0.1 ng Mg-28; T3: 60000 mGy/0.1 ng Mg-28; T2: 600000 mGy/0.1 ng Mg-28; and T1: 6000000 mGy/0.1 ng Mg-28. This signifies a delivery of concentrated radiation, rendering it possible to effectively eradicate cancer cells. This advantage of Mg-28 can be attributed to its identity as the Mg+2 ion, which serves as a cofactor in nearly 300 types of enzymes within cells. Given the rapid proliferation of cancer cells, their increased demand for magnesium ions in comparison to normal cells underscores the potential efficacy of this approach.

Further analysis reveals that the absorbed dose primarily results from the emission of beta particles. These particles exhibit a limited range of impact, spanning less than 3.6 mm in water. As a result, the surrounding healthy tissues adjacent to the tumor mass receive effective protection. This aspect highlights yet another remarkable advantage offered by Mg-28. Thus, enzymes reliant on magnesium ions as cofactors emerge as the prime targets for Mg-28 radioisotopes. Conversely, upon reaching their intended destination, these isotopes transform into agents capable of dismantling these very enzymes.

For stage 2A tumors, conforming to our previous assumption, they fall within the category of T4 tumor masses. Consequently, the utilization of Mg-28 can yield an impressively high absorbed dose, approximating 6000 mGy for a mere 0.1 ng of Mg-28. This dosage proves sufficiently potent to obliterate the entire cancerous mass. Even for larger tumors surpassing the confines of stage 2A, an option is to amplify the Mg-28 dosage by a factor of 10, resulting in an approximate value of 5.3 mCi or 197 MBq. This step is taken with the expectation of yielding improved outcomes. Nonetheless, it’s crucial to underscore that this viewpoint necessitates empirical validation through experiments.

## IV. Conclusion and Recommendations

The utilization of Mg-28, a radioisotope of the element magnesium, for the inactivation of magnesium-containing metalloenzymes stands as an innovative and well-founded concept. This approach underscores the potential application of nuclear transformation to disable metalloenzymes. Remarkably, metalloenzymes serve as precise targets for radioisotopes, which subsequently inactivate and irradiate them. The effectiveness of both actions holds the potential to inflict damage on tumor cells, should this method find application in cancer treatment research.

Radioactive Mg-28 ions possess the capability to displace stable Mg ions through competitive binding during metalloenzyme formation. With a competitive probability of 1‰, a mere 0.1 ng of Mg-28 can render approximately 2.5 billion magnesium-dependent metalloenzymes inactive. These cofactors play pivotal roles in numerous cellular metabolic processes, including DNA replication during the synthesis phase, RNA transcription across the G0, G1, and G2 phases, energy and glucose metabolism, as well as the initiation of cell cycle transitions. This strategy essentially serves as a form of chemotherapy, halting metabolic processes within tumor cells. The distinctiveness of this method lies in its reliance on a solitary radioisotope agent, Mg-28, which possesses the capacity to simultaneously or separately deactivate multiple magnesium-dependent metalloenzymes. This can be attributed to the fact that the decay of Mg-28 yields Al-28 (Al+3 ion) and subsequently Si-28 (Si+4 ion). Neither Al+3 nor Si+4 ions function as cofactors for magnesium-dependent enzymes. Consequently, their immediate inactivation leads to the disruption or hindrance of metabolic processes, profoundly affecting the cell cycle of cancer cells.

Armed with a mere 0.1 ng of Mg-28, its decay can generate remarkably high absorbed doses, ranging from 6000 to 6000000 mGy, specifically tailored for early-stage tumor masses with volumes ≤ 5 cc. This dose potency proves sufficient for the total elimination of the cancerous mass, while vital body tissues remain unscathed, as corroborated by MIRD calc simulations. The pivotal challenge lies in devising a means to transport Mg-28 to the tumor mass, minimizing the potential adverse effects on essential organs and neighboring healthy cells, either through radiation exposure or inactivation by Mg-28.

Hence, magnesium-dependent metalloenzymes find themselves as prime targets for Mg-28 radioisotopes. Conversely, the decay of this isotope results in the deactivation of these cofactors, all while concurrently delivering a robust radiation impact to the target with an exceptionally high probability and a substantial effective dose. This represents an advantage hitherto unparalleled by prior cancer treatment methods.

However, it’s crucial to acknowledge that this novel approach is currently presented in the form of hypothetical calculations and illustrative simulations. Validation through in vitro and in vivo experiments by scientists is imperative before embarking on clinical trials.

## Acknowledgments

This research was made possible through the support of the Ministry of Science and Technology of Vietnam under the basic research project with code: CS/21/02-03. The authors extend their gratitude to the Vietnam Atomic Energy Institute, the Nuclear Technology Center, and the Dalat Nuclear Research Institute for their invaluable assistance in facilitating the project.

## Contributions of the Authors

The primary author, Tran Van Luyen, conceived the research concept, carried out the principal investigations, assessed the outcomes, and authored this paper. Co-author Truong Hoang Tuan provided project oversight and facilitated the necessary procedures for its execution.

## Ethical Statement

This article adheres to ethical principles concerning research, experimentation, data analysis, and the dissemination of research findings.

## References

1. Enzym inactivation an overview in the book “Comprehensive Biotechnology” (Second Edition). Volume 1, 2011, Pages 25–39, https://doi.org/10.1016/B978-0-08-088504-9.00006-4

2. The enzym treatment of cancer and its scientific basic. New Spring Press 2009.

3. Amit K Jain 1, Sweta Jain, A C Rana. Metabolic enzym considerations in cancer therapy. Malays J Med Sci. 2007 Jan;14(1):10–7.

4. Robert D. Bruno and Vincent C.O. Nijar. Targeting Cytochrome P450 Enzyms: A New Approach in Anti-cancer Drug Development. Bioorg Med Chem. 2007 Aug 1; 15(15): 5047–5060.

5. Ge Yan Thomas Efferth. Chapter 11 - Identification of chemosensitizers by drug repurposing to enhance the efficacy of cancer therapy. Drug Repurposing in Cancer Therapy Approaches and Applications, 2020, Pages 295–310. https://doi.org/10.1016/B978-0-12-819668-7.00011-7.

6. https://www.brenda-enzyms.org/

7. Wilson JE. Hexokinases. Rev Physiol Biochem Pharmacol. 1995;126:65–198. [PubMed] [Google Scholar]

8. G. Maga,DNA Polymerases,Editor(s): Stanley Maloy, Kelly Hughes,Brenner’s Encyclopedia of Genetics (Second Edition),Academic Press,2013,Pages 376–378, https://doi.org/10.1016/B978-0-12-374984-0.00427-7.

9. Jeff Gelles and Robert Landick. RNA Polymerase as a Molecular Motor. Cell, Vol. 93, 13–16, April 3, 1998.

10. Abdullah, M; Al Alawi; Sandawana William Majoni and Henrik Falhammar: Magnesium and Human Health: Perspectives and Research Directions. Int J Endocrinol. 2018; 2018:9041694. doi: 10.1155/2018/9041694.

11. Fiorentini, D.; Cappadone, C.; Farruggia, G.; Prata, C. Magnesium: Biochemistry, Nutrition, Detection, and Social Impact of Diseases Linked to Its Deficiency. Nutrients 2021, 13, 1136. https://doi.org/10.3390/nu13041136.

12. Ebel, H.; Günther, T.; Günther, H.E.T. Magnesium Metabolism: A Review. Clin. Chem. Lab. Med. 1980, 18, 257–270. [Google Scholar] [CrossRef].

13. https://applets.kcvs.ca/IPTEI/pdf-elements/magnesium.pdf

14. Tran, L. (2019). A proposition for a cancer treatment study using radioactive metal co-factor enzyms. Biomedical Research and Therapy, 6(2), 2983–2985. https://doi.org/10.15419/bmrat.v6i2.519.

15. Luyen Van Tran. A Proposal for a Cancer Treatment Study Involving Radioactive Metal Co-factor Enzyms. Chapter 1. Print ISBN: 978-93-91473-87-7, eBook ISBN: 978-93-91473-90-7. http://DOI.10.9734/bpi/hmms/v15/9276D.

16. Tran Van Luyen. Deactivation DNA Polymerase and Hexokinase Enzyms by Radioisotope 28Mg in Cancer Treatment Research: An Advance Study. Chapter 2. Print ISBN: 978-93-91473-87-7, eBook ISBN: 978-93-91473-90-7. http://DOI.10.9734/bpi/hmms/v15/9277D.

17. Luyen TV. (2020) Deactivation DNA Polymerase and Hexokinase Enzyms by Radioisotope 28Mg in Cancer Treatment Research. BioMed Res J, 4(1): 196–201.

18. ICRP, 2012. Compendium of Dose Coefficients based on ICRP Publication 60. ICRP Publication 119. Ann. ICRP 41(Suppl.).

19. ICRP, 1988. Radiation Dose to Patients from Radiopharmaceuticals. ICRP Publication 53. Ann. ICRP 18 (1–4).

20. ICRP, 2007. The 2007 Recommendations of the International Commission on Radiological Protection. ICRP Publication 103. Ann. ICRP 37 (2–4).

21. https://mirdsoft.org/products/MIRDcalc_manual.pdf

22. ICRP, 2008. Nuclear Decay Data for Dosimetric Calculations. ICRP Publication 107. Ann. ICRP 38 (3).

23. https://blink.ucsd.edu/_files/safety-tab/rad/radionuclide/Mg-28.pdf.

24. https://www-nds.iaea.org/relnsd/vcharthtml/VChartHTML.html.

25. Takaaki Ogura, Natsuko I. Kobayashi, Hisashi Suzuki, Ren Iwata, Tomoko M. Nakanishi, Keitaro Tanoi; Magnesium uptake characteristics in Arabidopsis revealed by 28 Mg tracer studies. Planta, short communication. (2018)https://doi.org/10.1007/s00425-018-2936-4.

26. R. S. Becker and R. K. Sheline; Preliminary Experiments Using Mg-28 as a Tracer with Chlorophylls, Plants, and their Extracts;J. Chem. Phys. 21, 946 (1953); https://doi.org/10.1063/1.1699079.

27. Becker, Ralph S., and Raymond K. Sheline. “Biosynthesis and exchange experiments on some plant pigments using Mg28 and C14.” Archives of Biochemistry and Biophysics 54.2 (1955): 259–265.

28. Murdaugh, H. V., & Robinson, R. R. (1960). Magnesium excretion in the dog studied by stop-flow analysis. American Journal of Physiology-Legacy Content, 198(3), 571–574; doi:10.1152/ajplegacy.1960.198.3.571.

29. Care, A. D. (1960). The kinetics of magnesium distribution within the sheep. Research in Veterinary Science, 1(4), 338–349. Doi:10.1016/S0034-5288(18)34992-0.

30. Edwards Jr, H. M., Denis Nugara, and J. C. Driggers. “Fate of endogenous Mg28 in laying hens.” Poultry Science 41.6 (1962): 1975–1976.

31. Chutkow, Jerry G. “Studies on the metabolism of magnesium in the magnesium-deficient rat.” The Journal of Laboratory and Clinical Medicine 65.6 (1965): 912–926.

32. Gunther, T., J. Vormann, and V. Hollriegl. “effects of amiloride and furosemide on mg-28 transport into fetuses and maternal tissues of rats.” Magnesium-Bulletin 10.2 (1988): 34–37.

33. Lazzara, R., et al. “Tissue distribution, kinetics, and biologic half-life of Mg28 in the dog.” American Journal of Physiology-Legacy Content 204.6 (1963): 1086–1094.

34. Aikawa, Jerry K. “Gastrointestinal absorption of Mg28 in rabbits.” Proceedings of the Society for Experimental Biology and Medicine 100.2 (1959): 293–295.

35. McAleese, D. M., M. C. Bell, and R. M. Forbes. “Magnesium-28 studies in lambs.” The Journal of Nutrition 74.4 (1961): 505–514.

36. Field, A. C., and B. S. W. Smith. “Effect of magnesium deficiency on the uptake of 28Mg by the tissues in mature rats.” British Journal of Nutrition 18.1 (1964): 103–113.

37. Aikawa, Jerry K., et al. “Magnesium metabolism in rabbits using Mg28 as a tracer.” American Journal of Physiology-Legacy Content 197.1 (1959): 99–101.

38. Woodward, DAVID L., and DONAL J. Reed. “Uptake of 28 Mg and 45Ca by tissues of magnesium-deficient rabbits.” American Journal of Physiology-Legacy Content 217.5 (1969): 1483–1486.

39. Chutkow, Jerry G. “Metabolism of magnesium in the normal rat.” The Journal of Laboratory and Clinical Medicine 63.1 (1964): 80–99.

40. Wallach, Stanley, et al. “Tissue distribution and transport of electrolytes Mg28 and Ca47 in hypermagnesemia.” Metabolism 16.5 (1967): 451–464.

41. Aikawa, Jerry K., Gerald S. Gordon, and Eloise L. Rhoades. “Magnesium metabolism in human beings: studies with Mg28.” Journal of Applied Physiology 15.3 (1960): 503–507.

42. Avioli, L. V., and M. O. N. E. S. Berman. “Mg28 kinetics in man.” Journal of applied physiology 21.6 (1966): 1688–1694.

43. Aikawa, Jerry K., Eloise L. Rhoades, and Gerald S. Gordon. “Urinary and Fecal Excretion of Orally Administered Mg28.” Proceedings of the Society for Experimental Biology and Medicine 98.1 (1958): 29–31.

44. Kniffen, JARED C., et al. “Whole-body counter determination of 28Mg retention in humans.” Radioaktive Isotope in Klinik und Forschung. Urban and Schwarzenberg, Munchen, Berlin, Wein, 1970. 231–244.

45. Graham, Gastrointestinal. “absorption and excretion of Mg 28 in man.” Metabolism 9:p 646.

46. Watson, Walter S., et al. “Magnesium metabolism in blood and the whole body in man using 28 magnesium.” Metabolism 28.1 (1979): 90–95.

47. L. Silver, j. S. Robertson and l. K. Dahl; Magnesium turnover in the human studied with Mg-28. J Clin Invest. 1960;39(2):420–425. https://doi.org/10.1172/JCI104053.

48. Danielson, B. G. “Gastrointestinal magnesium absorption: Kinetic studies with 28Mg and a simple method for determination of fractional absorption.” (1979).

49. Anastassopoulou, J.; Theophanides, T. Magnesium-DNA interactions and the possible relation of magnesium to carcinogenesis. Irradiation and free radicals. Crit. Rev. Oncol. 2002, 42, 79–91. [Google Scholar] [CrossRef].

50. Yang, W. An overview of Y-family DNA polymerases and a case study of human DNA polymerase π. Biochemistry 2014, 53, 2793–2803. [Google Scholar] [CrossRef] [PubMed].

51. Lindahl, T.; Adams, A.; Fresco, J.R. Renaturation of transfer ribonucleic acids through site binding of magnesium. Proc. Natl. Acad. Sci. USA 1966, 55, 941–948. [Google Scholar] [CrossRef][Green Version]

52. Misra, V.K.; Draper, D.E. The linkage between magnesium binding and RNA folding 11 Edited by B. Honig. J. Mol. Biol. 2002, 317, 507–521. [Google Scholar] [CrossRef]

53. Tan, Z.J.; Chen, S.J. Importance of diffuse metal ion binding to RNA. Met. Ions Life Sci. 2011, 9, 101–124. [Google Scholar] [PubMed]

54. Fandilolu, P.M.; Kamble, A.S.; Dound, A.S.; Sonawane, K.D. Role of Wybutosine and Mg2+ Ions in Modulating the Structure and Function of tRNAPhe: A Molecular Dynamics Study. ACS Omega 2019, 4, 21327–21339. [Google Scholar] [CrossRef][Green Version]

55. Lange SS, Takata K, Wood RD (2011) DNA polymerases and cancer. Nat Rev Cancer 11:96–110.

56. Albertella MR, Lau A, O’Connor MJ (2005) The overexpression of specialized DNA polymerases in cancer. DNA Repair (Amst) 4:583–593.

57. Hoeijmakers JH (2009) DNA damage, aging and cancer. N Engl J Med 361: 1475–1485.

58. Principles and Practice of Biology Therapy of Cancer. Lippincott Williams & Wilkins: Philadelphia, PA, USA; 2000.

59. Baskar R, Lee KA, Yeo R, Yeoh KW (2012) Cancer and radiation therapy: Current advances and future directions. Int J Med Sci 9:193–199.

60. https://www.cancer.gov/about-cancer/treatment/types.

61. M.R.A. Pillai. Metallic Radionuclides and Therapeutic Radiopharmaceuticals. Dr Sc Thesis. Institute of Chemistry and Nuclear Technology. Wasawa, Poland. 2010

62. https://www.cancer.gov/types/prostate/research/radium-223-improves-survival.

63. Fallah J, et all. FDA Approval Summary: lutetium Lu 177 vipivotide tetraxetan for patients with metastatic castration-resistant prostate cancer. Clin Cancer Res. 2022 Dec 5:CCR–22-2875. doi: 10.1158/1078-0432.CCR-22-2875. PMID: 6469000.

64. https://www.cancer.org/cancer/thyroid-cancer/treating/radioactive-iodine.html.

65. Technical Reports Series No. 197. English STI/DOC/010/197 ¦ 92-0-115080-6

66. Christian, G.D. Medicine, trace elements, and atomic absorption spectroscopy. Anal. Chem. 1969, 41, 24A–40A. [Google Scholar] [CrossRef].

67. Planeta, K., Kubala-Kukus, A., Drozdz, A. et al. The assessment of the usability of selected instrumental techniques for the elemental analysis of biomedical samples. Sci Rep 11, 3704 (2021). https://doi.org/10.1038/s41598-021-82179-3

68. Al Alawi, A. M., Majoni, S. W., & Falhammar, H. (2018). Magnesium and Human Health: Perspectives and Research Directions. International Journal of Endocrinology, 2018. https://doi.org/10.1155/2018/9041694

69. Jahnen-Dechent, W., & Ketteler, M. (2012). Magnesium basics. Clinical Kidney Journal, 5(Suppl 1), i3. https://doi.org/10.1093/ndtplus/sfr163

70. T. J. Sprouse and D. E. Jolliffe, “Magnesium in cardiac tissue,” Archives of Biochemistry and Biophysics, vol. 171, no. 1, pp. 456–461, 1975.

71. J. M. Martin and J. B. Brown, “Magnesium content of normal human kidneys,” Clinical Science, vol. 26, no. 4, pp. 389–393, 1964.

72. A. Altura, B. Turlapaty, and B. Altura, “Magnesium, calcium, and other elements in organs of genetically hypertensive rats,” Hypertension, vol. 9, no. 6, pp. 631–636, 1987.

73. M. R. Levander, M. E. A. Johnson, and K. L. Wolf, “Magnesium content of human skeletal muscle,” Journal of the American College of Nutrition, vol. 13, no. 5, pp. 421–425, 1994.

74. S. Otsuka, Y. Iwasaki, and H. Kato, “Magnesium content in human pancreas and its alteration in diabetes,” Diabetes, vol. 35, no. 6, pp. 770–773, 1986.

75. T. M. Forrester, “Trace elements in human brain: Biopsy, necropsy, and in vivo studies,” in Trace Elements in Human Health and Disease, A. S. Prasad, Ed. New York: Academic Press, 1976, pp. 23–56.

76. P. H. Whanger and R. L. Weswig, “Magnesium and other minerals in human bone,” The Journal of Nutrition, vol. 96, no. 2, pp. 189–196, 1968.

